# Establishment and functional characterization of bovine endometrial epithelial organoids

**DOI:** 10.1101/2025.10.29.685416

**Authors:** Iebu Devkota, Zachary L. Bonomo, Dailin M. Fuego, Yuxia Li, Xujia Zhang, Shavahn C. Loux, Charles R. Looney, Ana I. V. Maia, Fabrizio Donnarumma, Antonios Matsakas, Anastasios Vourekas, Philip H. Elzer, Xing Fu, Kenneth R. Bondioli, John J. Bromfield, Pablo Bermejo-Àlvarez, Constantine A. Simintiras

## Abstract

Pre-implantation embryonic loss constitutes a major barrier to reproductive efficiency in livestock, yet the extrinsic determinants of embryonic survival remain poorly defined. Intra-organoid fluid (**IOF**) faithfully recapitulates native tissue secretions across multiple organ systems. We hypothesized that bovine endometrial epithelial organoids (**BEEO**) would produce IOF that mirrored *in vivo* uterine luminal fluid composition and extend embryo culture duration *in vitro*. We pursued three objectives: (***a***) establish and morphologically characterize BEEO, (***b***) define BEEO transcriptomic and secretory responses to estradiol (**E2**), medroxyprogesterone acetate (**MPA**), and interferon-tau (**IFNτ**), and (***c***) determine whether BEEO-derived IOF can support *in vitro* embryonic development beyond Day 8 (hatched blastocyst stage) under conventional culture conditions. BEEO were established from primary endometrial tissue (n=4) and maintained a stable epithelial phenotype through multiple passages. Transcriptomic profiling revealed robust responses to stimulation, with E2, MPA, and IFNτ inducing distinct gene expression programs consistent with *in vivo* effects. IOF metabolomic analysis confirmed hormone-dependent regulation of IOF secretory output, with E2+MPA (diestrus mimic) enhancing the production of metabolites implicated in conceptus development. Remarkably, IOF from diestrus mimic-stimulated BEEO, despite being diluted approximately seven-fold in PBS, maintained embryo survival rates comparable to optimized commercial medium, and exceeded PBS-only controls. These findings position BEEO as a physiologically relevant model for dissecting maternal-embryo interactions *in vitro* and identifying targets to improve fertility in cattle and other livestock.

## INTRODUCTION

Developmental failure during early pregnancy is a major bottleneck to livestock reproductive efficiency and imposes economic burdens via prolonged calving intervals, decreased milk yield, slowed genetic gain, and repeated insemination costs (De Vries, 2006; Perkel *et al*., 2015). Approximately 55% of bovine pregnancy losses occur prior to Day 21, coinciding with critical developmental processes including conceptus (embryo and extra-embryonic membranes) elongation, maternal pregnancy recognition, and implantation (Walsh *et al*., 2011; Wiltbank *et al*., 2016). Conceptus elongation represents a particularly vulnerable period of pregnancy loss, with one-third to one-half of embryos failing to survive this transition (Dunne *et al*., 2000; Santos *et al*., 2004; Berg *et al*., 2010; Crowe *et al*., 2025).

Under physiological conditions, the elongating conceptus undergoes sequential morphological transformations from a spherical (Day 8) ovoid (Day 12), tubular (Day 14), and finally a filamentous (Day 16) structure (Betteridge *et al*., 1980; Simintiras *et al*., 2021a). Conceptus elongation is fundamental, as the concentration of the conceptus-derived pregnancy recognition signal [interferon tau (**IFN𝛕**) (Bazer and Thatcher, 2017)] produced is proportional to conceptus size (Kerbler *et al*., 1997). Thus, inadequate conceptus elongation results in insufficient IFN𝜏 to trigger an appropriate maternal response, resulting in pregnancy loss (Sánchez *et al*., 2019).

Studying the determinants of conceptus survival is constrained by current conventional *in vitro* culture systems which cannot support embryo development much beyond the hatched blastocyst stage (approximately Day 8). Despite ongoing efforts, conceptus elongation has not been successfully recapitulated *in vitro* (Maddox-Hyttel *et al*., 2003; Brandão *et al*., 2004; Vajta *et al*., 2004; Alexopoulos *et al*., 2006; Zhao *et al*., 2015). Partial progress has been achieved by extending bovine embryo culture to Day 12 through culture medium modifications (Ramos-Ibeas *et al*., 2020, 2022, 2023). *In vivo*, conceptus elongation fails in the absence of uterine glands (Gray *et al*., 2001), further underscoring the dependence of this process on uterine luminal fluid – a hormonally regulated, dynamic mixture of proteins, metabolites, and other biochemicals (Forde *et al*., 2014; Simintiras *et al*., 2019a–c, 2022).

Organoids – self-organizing, three-dimensional cell culture systems (Zhao *et al*., 2022) – represent a powerful platform for investigating tissue-specific interstitial fluid composition regulation and function *in vitro*. Recent advances include: (***a***) snake venom gland organoids produce biologically active venom (Post *et al*., 2020), (***b***) human lacrimal gland organoids generate tears (Bannier-Hélaouët *et al*., 2021), (***c***) human choroid plexus organoids produce cerebrospinal fluid (Pellegrini *et al*., 2020), and (***d***) human endometrial epithelial organoids secrete a uterine-like intra-organoid fluid (**IOF**) (Simintiras *et al*., 2021b). Despite these advances, to our knowledge, just two studies have reported bovine endometrial epithelial organoid (**BEEO**) culture (Nishino *et al*., 2021; Kwon *et al*., 2025), providing important initial groundwork and representing a significant gap in livestock reproductive biology research tools.

Here, we establish and functionally characterize BEEO to probe epithelial function and maternal-conceptus crosstalk. Our specific objectives were to: (***a***) establish and morphologically characterize BEEO, (***b***) define BEEO transcriptomic and secretory responses to estradiol (**E2**), a synthetic progesterone (**P4**) analogue medroxyprogesterone acetate (**MPA**) (Bläuer *et al*., 2005), and interferon-tau (**IFNτ**), and (***c***) determine if BEEO-derived IOF can support *in vitro* embryonic development beyond the current limit of approximately Day 8 under otherwise conventional culture conditions.

## MATERIALS AND METHODS

### Animals

All animal-related procedures were approved by the Louisiana State University Agricultural Center Institutional Animal Care and Use Committee protocols. The estrous cycles of four crossbred cattle, with a mean (± SD) age of 10.8 ± 3.3 years and weight of 443.3 ± 28.5 kg, were synchronized using an established protocol (Simintiras *et al*., 2018), involving intramuscular administration of a gonadotropin-releasing hormone (**GnRH**) analogue (Fertagyl; Merck Animal Health, 014148) alongside intravaginal insertion of a P4 (1.38 g) controlled internal drug release (**CIDR**) device (Eazi-breed; Zoetis Animal Health, 021274). Seven days later, prostaglandin F2α (**PGF2α**) (Lutalyse; Zoetis Animal Health, 026256) was administered intramuscularly, followed by CIDR removal the following day. Estrus detection was performed using breeding indicators (Estrotect; Rockway, 78013). Indicators were monitored twice daily starting 48 h following CIDR withdrawal. All animals displayed signs of standing estrus and were sacrificed on Day 5 of the estrous cycle. This is summarized in **Supplementary Figure 1A**.

### Sample generation

Complete reproductive tracts from synchronized cattle (n=4) were obtained at a local commercial abattoir (Coastal Plains Meat Company, Eunice, LA), immediately placed in sealed plastic bags on ice, and transported to the laboratory within 1.5 h. Upon arrival, the presence of a stage-appropriate ovarian *corpus luteum* was confirmed, and the corresponding ipsilateral uterine horn was dissected along the anti-mesometrial axis. Intra-caruncular endometrial biopsies (*i*.*e*., comprising both luminal and glandular epithelia) were obtained and either (***a***) placed in 4 % (*v*/*v*) formaldehyde (VWR, 10790-712) in Milli-Q [18.2 MΩ·cm] water and kept at 4 °C until immunohistochemical analysis, or (***b***) processed for BEEO establishment. The complete workflow is summarized in **Supplementary Figure 1B**.

### Initial organoid establishment

To generate BEEO, biopsies were finely minced and enzymatically digested in Dulbecco’s Modified Eagle Medium (**DMEM**)-F12 medium (Gibco-Invitrogen, 11320033) comprising 1 mg·ml^−1^ Collagenase V (Sigma-Aldrich, 9001-12-1) and 1.2 mg·ml^-1^ Dispase II (Sigma Aldrich, D4693) – at 38.5 °C in 5 % CO_2_ and 5 % O_2_ under N_2_ for 1.5 h with vigorous agitation (ThermoScientific, 88881103B). Digestion was terminated by adding an equal volume of *Neutralizing Medium*. All media compositions are provided in **Supplementary Table 1**.

Undigested fragments were allowed to sedimented by gravity for approximately 10 min before supernatant collection and 70 µm filtration (Corning, 431751). The filtrate was centrifuged (300 × *g*, 10 min, 4°C) before pellet resuspension in chilled (4 °C) *Wash Medium*. Following re-centrifugation (300 × *g*, 10 min, 4°C) the pellet was resuspended in pre-equilibrated (38.5 °C) *Stromal Medium*. Cell suspensions were counted by trypan blue (VWR, K940) exclusion using an automated cell counter (Corning, 6749), and seeded to 75 cm^2^ flasks (VWR, 10062-860) at a density of 2 × 10^6^ per flask. Cells were cultured at 38.5 °C under 5 % CO_2_ in humidified air for 18 h to achieve stromal, but not epithelial, cell attachment (Turner *et al*., 2014).

Non-adhered epithelial cells were collected, pelleted (300 × *g*, 10 min, 4°C), and resuspended in chilled (4 °C) *Wash Medium*. The cell suspension was counted as above, re-centrifuged (300 × *g*, 10 min, 4°C), and resuspended in chilled (4 °C) Cultrex hydrogel (R&D Systems, 3433-005-R1) at a volume achieving a dilution of 500 cells·µl^-1^. Working on ice, cells were transferred to 12 well plates (Fisherbrand, FB012928) at a density of five 20 µl hydrogel domes per well, before incubation at 38.5 °C under 5 % CO_2_ in humidified air for 30 min to facilitate hydrogel polymerization. Each well was then flooded with pre-equilibrated (38.5 °C) *Thawing Medium*.

### Cryopreservation

To evaluate capacity for cryopreservation, BEEO were first removed from the surrounding hydrogel by replacing conditioned *Expansion Medium* with chilled (4 °C) *Wash Medium* before manually dislodging the hydrogel using a wide-orifice 1000 µl pipette tip (Thermo Scientific, 02707008). The cell-hydrogel suspension was transferred to a conical tube (Falcon, 352196), pelleted (300 × *g*, 10 min, 4°C), and resuspended in chilled *Wash Medium*. This wash step was repeated until the hydrogel was no longer visible by eye. The resulting pellet was resuspended in *Freezing Medium* and stored at −80 °C within a freezing container (Thermo Scientific, 5100-0036), to achieve a cooling rate of approximately 1 °C·min^-1^.

The following morning, cells were stored in N_2_(*l*) for roughly 3 months, after which BEEO were rapidly thawed by vial immersion in a pre-equilibrated (38.5 °C) water bath for 45 sec. The suspension was then immediately diluted in chilled *Wash Medium* within a 15 ml conical tube. centrifuged (300 × *g*, 10 min, 4 °C) and resuspended in chilled *Expansion Medium*. Thereafter, cell counting, centrifugation, Cultrex hydrogel resuspension, plating, and incubation in *Expansion Medium*, was performed as aforementioned.

### Sub-culture

After cryopreservation and thawing, BEEO were maintained at 38.5 °C under 5 % CO_2_ in humidified air with medium replenishment every 48 h for 6 days, by which point BEEO began to form. More specifically, *Thawing Medium* was used for the first two media changes, after which *Expansion Medium* was used for all subsequent media changes. BEEO were monitored daily and sub-cultured at an approximate split ratio of 1:3 every 10-15 days, based on visual confluency determination. All experiments were conducted at the third subculture.

### Hormonal supplementation

Following an initial 6-day culture expansion period, BEEO from each biological replicate (n=4) were randomly allocated to one of six experimental groups (**Fig**. **1A**). All groups received *Base Medium* throughout the treatment period. Hormonal treatments consisted of (***a***) 10 nM E2 (MP Biomedicals, 194565) dissolved in ethanol yielding a final solvent contribution of 0.33% (*v*/*v*), (***b***) 1 µM MPA (Acros Organics, 461120010) dissolved in ethanol yielding a final solvent contribution of 0.33% (*v*/*v*), and/or (***c***) 5 nM ovine IFNτ (Bio world, 22060609-1) dissolved in water (18.2 MΩ·cm). E2, MPA, and IFNτ concentrations were guided by analogous human endometrial organoid (Turco *et al*., 2017) and ovine embryo *in vitro* supplementation (Wang *et al*., 2013) literature. The vehicle control group was supplemented with 0.67% (*v*/*v*) ethanol, corresponding to the highest solvent concentration among treatment groups. Media was replenished every 24 h during the treatment period, defined as Day 0 onwards.

**Figure 1.**
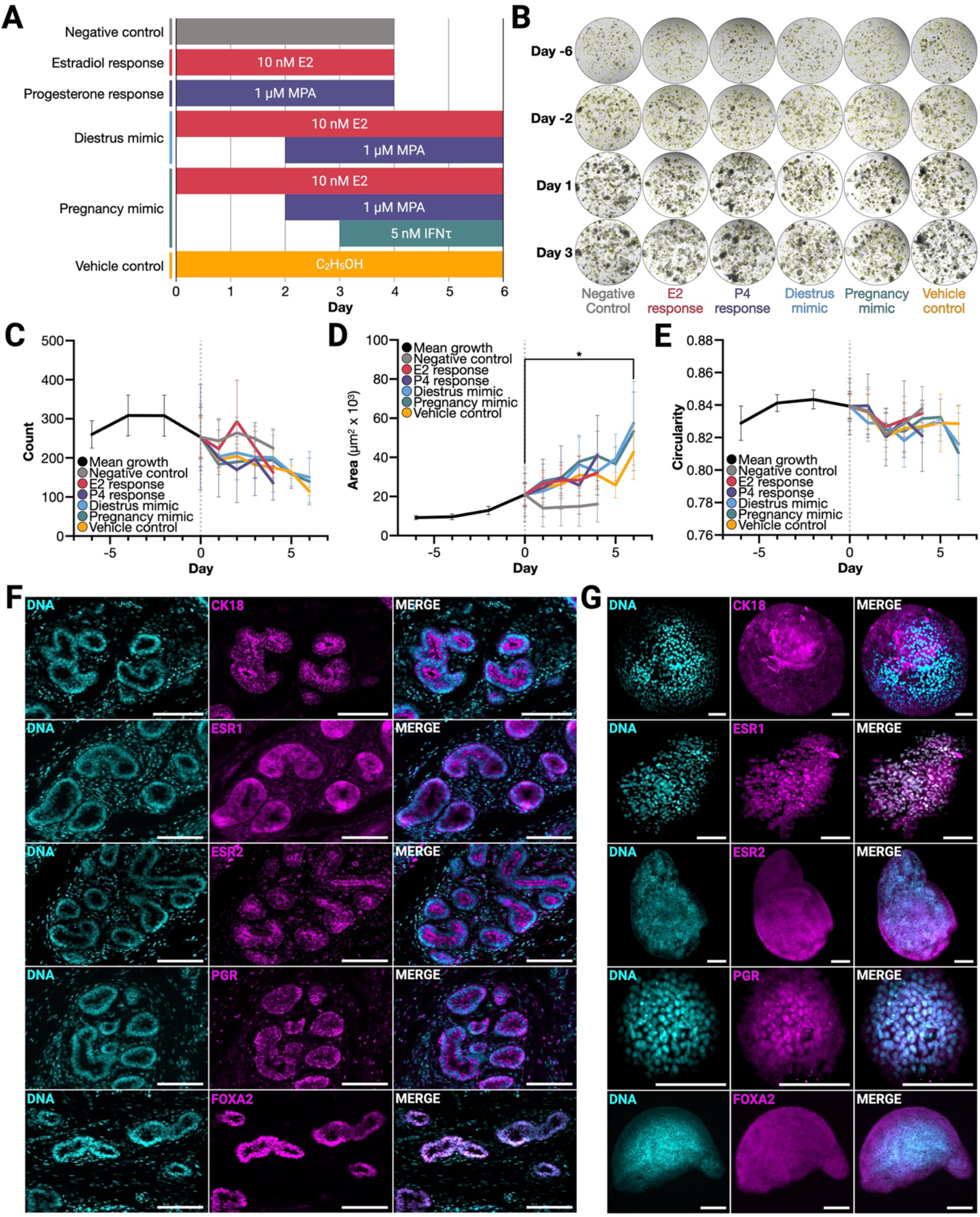
Morphological characterization of bovine endometrial epithelial organoids (BEEO). (**A**) Dependent experimental variables – BEEO were treated with 17β-estradiol (E2), medroxyprogesterone acetate (MPA), interferon tau (IFNτ), ethanol (C2H5OH), or not (negative control) for 96 or 144 h. (**B**) Representative organoid brightfield images across treatments and time. Note: Yellow borders are software generated. (**C**-**E**) Mean (±SEM) BEEO (n=4) morphometric kinetics – (**C**) count, (**D**) cross-sectional area, and (**E**) circularity. Effect of time [*P*≤0.05 (*)] but not treatment observed. (**F**) Representative *ex vivo* endometrial immunohistochemistry. (**G**) Representative BEEO immunohistochemistry following E2 treatment. All scale bars: 100 µm. Additional abbreviations: progesterone (P4), cytokeratin 18 (CK18), estrogen receptor alpha (ESR1), estrogen receptor beta (ESR2), progesterone receptor (PGR), and forkhead box A2 (FOXA2).

### Morphological assessment

BEEO were monitored by brightfield microscopy at 5× magnification (Leica DMI8) every 48 h throughout the 6-day expansion phase and every 24 h during treatment. One hydrogel dome per image was randomly selected for quantitative morphometric analysis. Representative images are provided in **Figure 1B**. Using Fiji (ImageJ, National Institutes of Health) with the SAMJ plugin, all BEEO in each image were manually selected via the *Points* function within the EfficientViTSAM-I2 module. Individual BEEO count, circularity, and area data were recorded.

To facilitate meaningful inter-group comparisons and account for baseline variability between groups, morphometric parameters [count (**Fig**. **1C**), circularity (**Fig**. **1D**), and area (**Fig**. **1E**)] were normalized to a common starting point. More specifically, normalization was achieved by calculating the difference between each group mean measurement at Day 0 and the negative control group reference value, before applying this offset as a constant adjustment factor to all subsequent time points within that treatment group. This preserved the relative changes and dynamics within each group while enabling direct statistical comparison of treatment effects by eliminating differences in initial BEEO trajectories. Data analysis (means ± SEM) was performed in Microsoft Excel for Mac (v. 16.97.2, b. 25052611). Graphs were generated in GraphPad Prism for Mac (v. 10.4.2, b. 534). Statistical procedures are detailed below.

### Tissue immunohistochemical labelling

Endometrial biopsies were fixed by immersion in 4% (*v*/*v*) aqueous formaldehyde (VWR, 10790-712) in Milli-Q (18.2 MΩ·cm) water at 4 °C for 72 hours. Tissues were then rinsed by serial immersion in 1× phosphate-buffered saline (**PBS**; Fisher Bioreagents, BP39920) and trimmed to approximately 1 cm^2^. Samples were then immersed in 30% (*v*/*v*) sucrose (Fisher Bioreagents, BP220-212) in 1× PBS at 4 °C overnight. Tissues were subsequently equilibrated at room temperature (**RT**) for 45 min in a 1:1 mixture of 30% (*v*/*v*) sucrose in PBS and Tissue-Tek Optimal Cutting Temperature (**OCT**) compound (Finetek, 4583), then embedded in 1× OCT compound within cryomolds (Finetek, 4566), and stored at −80 °C. Embedded tissue blocks were cryo-sectioned (Thermo Scientific, HM525NX) at a thickness of 5 µm, mounted onto charged slides (VWR, 48311-703), and stored at −20 °C.

Prior to immunohistochemistry (**IHC**), mounted sections were incubated at 37 °C for 5 min and post-fixed in 4% (*v*/*v*) aqueous formaldehyde for 10 min at RT with orbital shaking. Slides then underwent three sequential 1× PBS washes (10 min each) followed by two sequential water (18.2 MΩ·cm) rinses (2 min each). Antigen retrieval was performed by slide incubation in pre-heated 10% (*v*/*v*) Reveal Decloaker solution (BioCare Medical, RV1000MMRTU) at 98 °C for 30 min. Slides were allowed to return to RT for 1 h before two sequential 1× PBS washes (5 min each) with orbital shaking. Tissue sections were encircled using a hydrophobic barrier pen (Electron Microscopy Sciences, 71310) and blocked by the addition of 10% (*v*/*v*) Normal Goat Serum (NGS; AbCam, ab7481) in 1× PBS for 1 h, at RT within a humidified light-protected chamber (Electron Microscopy Sciences, 71397-B).

Primary antibodies (**Supplementary Table 2**), diluted 1:300 in 10% (*v*/*v*) NGS in 1× PBS, were applied overnight at 4°C in a humidified light-protected chamber. After three immersions in 1× PBS with orbital shaking (10 min each), the secondary antibody (**Supplementary Table 2**), diluted in 1:400 in 10% (*v*/*v*) NGS in 1× PBS, was applied for 1 h, at RT. Following an additional three washes in 1× PBS, nuclear counterstaining was performed using Hoechst 34580 (AAT Bioquest, 17537), diluted 1:5000 in PBS, and applied to slides for 5 min at RT, followed by three PBS washes (5 min each). Finally, sections were mounted with Fluoromount-G (Invitrogen, 00-4958-02) and covered (VWR, 48404-453). Imaging was performed using a Leica DMI8 fluorescence microscope coupled to Leica Application Suite X (LASX) software (v. 3.9.128433). Representative images are provided in **Figure 1F**.

### Whole-mount BEEO immunohistochemical labelling

Following treatment, select culture wells were designated for immunofluorescence imaging. To achieve this, *Expansion Medium* was replaced with chilled (4 °C) *Wash Medium* before manual hydrogel dislodgement as aforementioned. The cell-hydrogel suspension was transferred to a conical tube, centrifuged (300 × *g*, 4 min, 4°C), and resuspended in chilled *Wash Medium*. This wash step was repeated until the hydrogel was no longer visible by eye.

BEEO fixation was achieved by pellet resuspension and incubation in 4% (*v*/*v*) aqueous formaldehyde in water (18.2 MΩ·cm) for 20 min at RT with orbital shaking. BEEO were then centrifuged (300 × *g*, 10 min, 4°C) and resuspended in 1× PBS. This step was repeated three times before BEEO storage at 4 °C for no longer than one week. Subsequent permeabilization was achieved by BEEO centrifugation (300 × *g*, 10 min, 4°C), resuspension and incubation in 1× PBS + 0.1% (*v*/*v*) Triton X-100 (Apacor, 1499) for 30 min, at RT with orbital shaking. Post-centrifugation (300 × *g*, 10 min, 4°C), BEEO were resuspended and incubated in 1× PBS + 5% (*w*/*v*) bovine serum albumin (**BSA**; Sigma Aldrich, A6003) for 1 h, at RT with orbital shaking.

Thereafter, BEEO were centrifuged (300 × *g*, 4 min, 4 °C) before resuspension and overnight incubation at 4 °C with the primary antibody (**Supplementary Table 1**), diluted to 1 µg·ml^-1^ in *Antibody Buffer* [1.0 % (*v*/*v*) BSA and 0.1 % (*v*/*v*) Tween-20 (Sigma-Aldrich, P7949) in 1× PBS]. BEEO were washed three times – each wash involving centrifugation (300 × *g*, 4 min, 4 °C) and resuspension in 0.1 % (*v*/*v*) BSA and 0.1 % (*v*/*v*) Tween-20 (Sigma-Aldrich, P7949) in 1× PBS – before incubation for 1 h, at RT with the secondary antibody (**Supplementary Table 1**) diluted to 1 µg·ml^-1^ in *Antibody Buffer*. After two further PBS washes, BEEO were resuspended and incubated in Hoechst (1:5000 in 1× PBS) for 10 min, at RT with orbital shaking, washed three more times with PBS, and transferred onto slides using an EZ-Grip (Cooper Surgical, 7722802). Finally, BEEO were mounted in Fluoromount-G, covered, and imaged as aforementioned. Representative images are provided in **Figure 1G**.

### RNA extraction

Immediately following IOF isolation (described below), RNA was extracted from each residual BEEO pellet using the PureLink RNA Mini Kit (Invitrogen, 12183018A) in accordance with manufacturer instructions. In brief, epithelia were lysed by addition of *Lysis Buffer* supplemented with 2-mercaptoethanol (Millipore-Sigma, 444203250ML). The lysate was then homogenized by vortex and an equal volume of aqueous 70 % (*v*/*v*) ethanol (Koptec, V1001) was added to each lysate. Mixtures were transferred into spin cartridges and centrifuged (12,000 × *g*, 15 sec, RT). The flow-through was discarded, and cartridges washed with *Wash Buffer I* once and *Wash Buffer II* twice, with centrifugation as above following each wash. After the third wash, spin cartridges were centrifuged (12,000 × *g*, 2 min, RT) to remove residua buffer. RNA was eluted by addition of 30 μl DEPC-treated RNase free water (Thermo Scientific, R0601) to each cartridge before incubation for 1 min at RT. Each column was then centrifuged (12,000 × *g*, 2 min, RT) for RNA collection. RNA yield and purity were determined using a NanoDrop One spectrophotometer). RNA was stored at −80 °C until analysis. RNA concentrations recovered from each sample are provided in **Supplementary Table 3**.

### cDNA library preparation and RNA-sequencing

A total of 200 ng RNA per sample was processed using a low input RNA library preparation kit (New England Biolabs, E6420L) in line with manufacturer instructions. In brief, reverse transcription was performed using a template-switching mechanism to synthesize full-length cDNA, followed by amplification with 8 polymerase chain reaction (**PCR**) cycles to yield sufficient material. Amplified cDNA was purified using solid-phase reversible immobilization (SPRI) Select beads (Redoxica, DN9004), before fragmentation and end-repair using a fragmentation system kit (New England Biolabs, E7805L), also in accordance with manufacturer instructions. Adapters were ligated, and cDNA libraries were enriched via 8 PCR cycles using multiplex oligonucleotides for dual index primers (New England Biolabs, E6446S). Libraries were quantified using a Qubit 4 (Thermo Fisher Scientific, Q33238) fluorometric assay and assessed for size distribution and quality using an Agilent Bioanalyzer (Agilent Technologies) with corresponding buffers (Agilent Technologies, 5067-5589) and screen tape (Agilent Technologies, 5067-5588). Libraries were stored at −20 °C until shipment for paired-end sequencing (150 bp reads; PE150) on the Illumina NovaSeq X Plus platform at Novogene.

### Transcriptomic analysis

Raw FASTQ reads were adapter and quality trimmed using TrimGalore (v. 0.6.10), then mapped to the *Bos taurus* genome (ARS-UCD1.3) using STAR (v. 2.7.11b). Settings to generate raw read counts were as follows (--outSAMstrandField intronMotif --outFilterIntronMotifs RemoveNoncanonicalUnannotated --alignEndsType Local --chimOutType WithinBAM --twopassMode Basic --twopass1readsN -1 --quantMode GeneCounts), with reads annotated to ENSEMBL 113. Normalization and differential expression analysis of raw RNA-seq counts (**Supplementary Table 4**) was performed in EdgeR (v. 4.0.16). Genes with at least 10 counts in at least 20% of samples were retained, normalized using the gene length corrected trimmed mean of M-values method (Smid *et al*., 2018), and modeled with a design matrix incorporating treatment groups and biological replicates (batch correction). Dispersion estimation and quasi-likelihood F-testing identified differentially expressed genes (**DEG**) as those with a Benjamini– Hochberg false discovery rate (FDR) ≤ 0.05 and an absolute log2 fold change (FC) > 1.

Principal component analysis (**PCA**) was conducted using SVA (v. 3.50.0) *ComBat* batch-corrected logarithmically transformed counts per million (**CPM**) values, preserving treatment effects via a design matrix. PCA was performed using *prcomp* with unit variance scaling – the first two components were visualized using *ggplot2* (v. 3.5.2) with 95% confidence ellipses and variance explained on the axes (**Fig**. **2A**). Co-regulated DEG clusters were identified by k-means clustering (k = 6) of the 2,410 most variable genes (z-score scaled, 25 random starts). A heatmap (**Fig**. **2B**) was generated using *pheatmap* (v. 1.0.12), ordering genes by hierarchical clustering of cluster means and samples by treatment progression.

**Figure 2.**
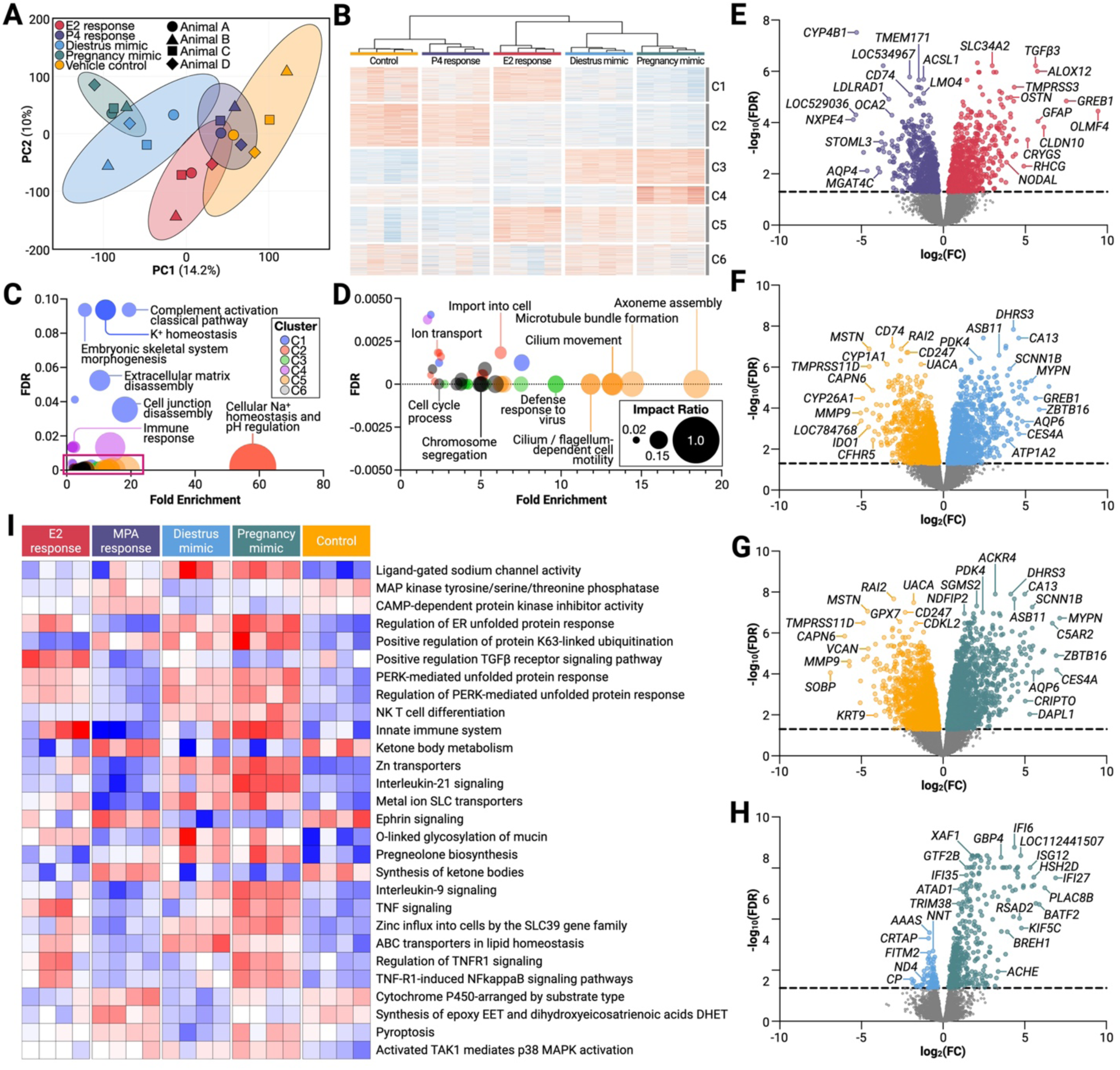
Bovine endometrial epithelial organoid (BEEO) transcriptomic characterization. (**A**) Principal component analysis of BEEO transcriptomic profiles. (**B**) Heatmap with hierarchical clustering of top differentially expressed genes (DEG) by BEEO treatment. (**C**-**D**) Multivariate plot of DEG pathway significance and impact according to cluster. Bubble size is proportional to the impact ratio (number of genes divided by total pathway size), whereas the vertical axis unit is inverse false discovery rate (FDR). Note that panel (D) provides a magnified view of the red highlighted area within panel (C). (**E**-**H**) Volcano plots of DEG between BEEO treated with: (**E**) 17β-estradiol (E2) *vs*. medroxyprogesterone acetate (MPA). Red and blue indicate E2 and MPA upregulation, respectively, (**F**) diestrus mimic *vs*. vehicle control. Blue and yellow indicate diestrus mimic and control upregulation, respectively, (**G**) pregnancy mimic *vs*. vehicle control. Green and yellow indicate pregnancy mimic and control upregulation, respectively, and (**H**) pregnancy mimic *vs*. diestrus mimic. Green and blue indicate pregnancy mimic and diestrus mimic upregulation, respectively. (**I**) Parametric gene set enrichment analysis using gene ontology pathway annotations, highlighting activated (red) and suppressed (blue) processes in BEEO across treatments. Additional abbreviations: clusters 1-6 (C1-C6).

Cluster-associated gene ontology (**GO**) terms, specifically *biological process* and *molecular function*, were identified using iDEP (v. 2.0.1) and visualized as bubble plots using GraphPad Prism (v. 10.4.2, b.534), with bubble diameter representing the impact ratio – the proportion of DEG in a pathway relative to the total number of genes annotated to that pathway (**Fig**. **2C**-**D**). DEG volcano plots (**Fig**. **2E**-**H**) of pairwise treatment comparisons were generated in GraphPad Prism from EdgeR derived results. Finally, Parametric Gene Set Enrichment Analysis (**PGSEA**) was conducted using the iDEP Pathway module, applying an FDR threshold of 0.1 to GO *biological process* and *molecular function* databases (**Fig**. **2I**).

### Intra-organoid fluid extraction

IOF isolation followed an established centrifugation method (Simintiras *et al*., 2021b). Specifically, BEEO were removed from the surrounding hydrogel by replacing conditioned *Expansion Medium* with chilled (4 °C) *Wash Medium* before manually dislodging the hydrogel as aforementioned. The cell-hydrogel suspension was transferred to a conical tube, centrifuged (300 × *g*, 10 min, 4°C), and resuspended in chilled *Wash Medium*. This wash step was repeated until the hydrogel was no longer visible by eye. Two additional washes using 1× PBS yielded clear supernatants, which were discarded. BEEO were then resuspended in 100 µl 1× PBS, transferred to 1.5 ml tubes (VWR, 89000-028), and centrifuged (4,000 × *g*, 30 min, 4 °C) to release IOF within the supernatant. Supernatant volumes were recorded before being snap-frozen in N_2_(*l*), stored at −80°C (overnight), and preserved in N_2_(*l*) until further analysis.

To account for potential differences in BEEO mass, IOF volumes were calculated as the ratio of supernatant volume (µl) to total RNA concentration (ng·µl^-1^) from residual pellets. This yields units of µl^2^·ng^-1^ with results expressed as the mean ± standard error of the mean (**SEM**) for four biological replicates (**Fig**. **3A**). Thereafter, IOF from each group was pooled, and total protein concentration (mg·ml^-1^) was quantified using a NanoDrop One spectrophotometer (Thermo Fisher Scientific). As such, protein measurements were not normalized to total RNA, and results are presented as the mean ± SEM from technical triplicates (**Fig**. **3B**).

**Figure 3.**
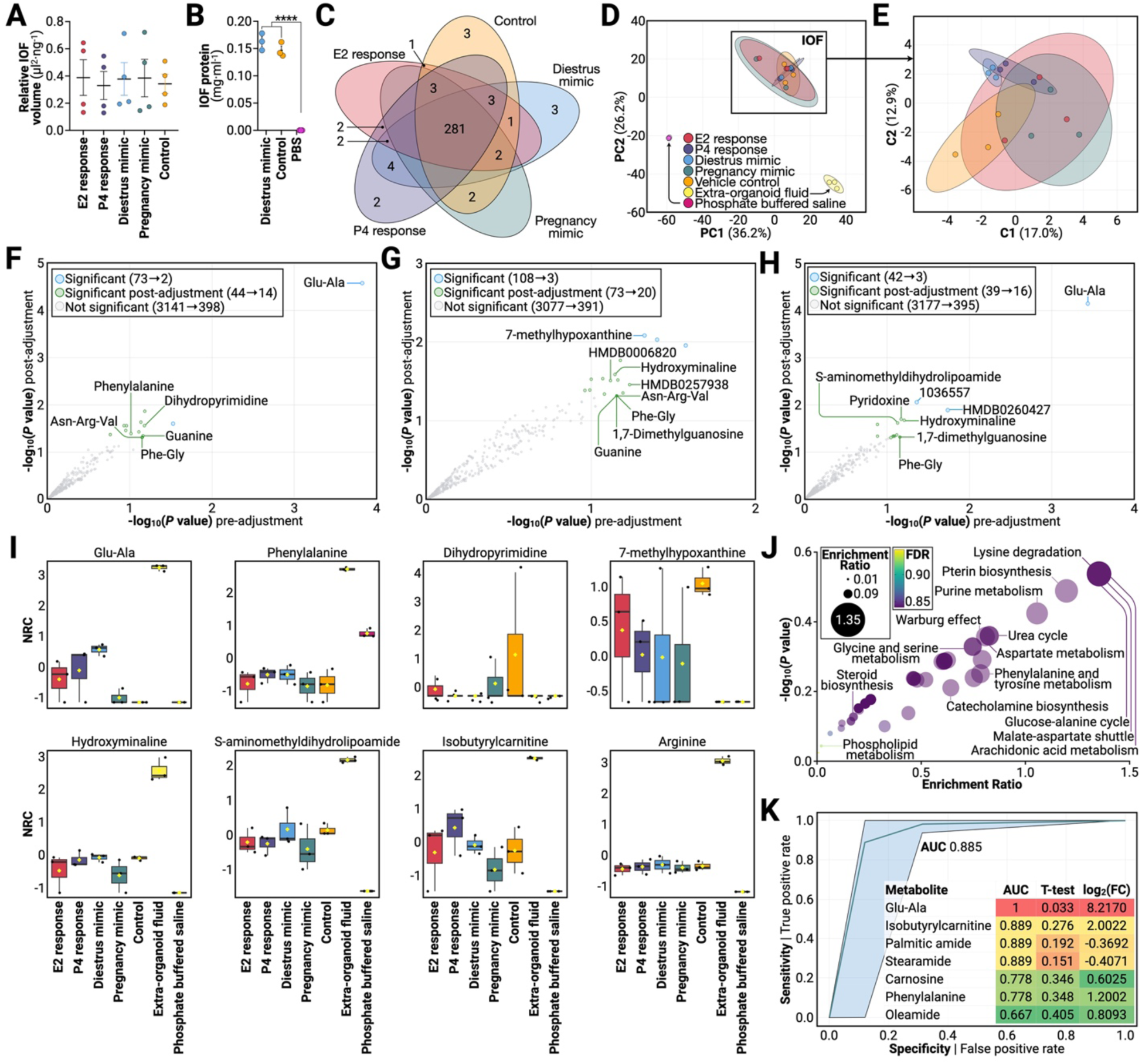
Bovine endometrial epithelial organoid (BEEO) intra-organoid fluid (IOF) metabolomic characterization. (**A**) Normalized IOF volume harvested by treatment. (**B**) Total protein present in IOF from diestrus mimic and vehicle control treated BEEO in addition to phosphate buffered saline (PBS) control [*P*≤0.0001 (****)]. (**C**) Number of unique metabolites detected in IOF across BEEO treatments. (**D**) Principal component analysis of treatment-specific IOF, extra-organoid fluid, and PBS control. (**E**) Sparse partial least squares discriminant analysis of IOF metabolomes by BEEO treatment. (**F**-**G**) Linear models with covariate (animal batch) adjustment comparing IOF metabolomic profiles between (**F**) diestrus mimic *vs*. vehicle control, (**G**), pregnancy mimic *vs*. vehicle control, and (**H**) diestrus *vs*. pregnancy mimic treated BEEO. (**I**) Boxplots of select metabolite normalized relative concentrations (NRC) across treatments. Central horizontal lines represent the median value with outer box boundaries depicting upper and lower quartile limits. Error bars depict the minimum and maximum distributions, with yellow rhombi representing the mean values. (**J**) Pathway enrichment analysis using differentially abundant IOF metabolites from diestrus *vs*. pregnancy mimic treated BEEO. (**K**) Receiver operating characteristic (ROC) curves generated by input of 7 metabolites (insert) to predict IOF metabolite biomarkers between diestrus *vs*. pregnancy mimic treated BEEO. Corresponding individual area under the ROC curve (AUC), t-test *P* values, and fold change (FC) values also provided.

### Mass spectrometry

High-throughput, untargeted metabolomic profiling of IOF was performed at the Louisiana State University Mass Spectrometry Facility. Prior to analysis, IOF was thawed on ice for 60 min. Thereafter 50 µl of each sample was supplemented with 100 µl methanol and 100 µl acetonitrile. Tubes were vortexed for 15 sec, sonicated for 5 min, and incubated at −20 °C for 1 h, followed by centrifugation (15,000 × *g*, 10 min, 4 °C).

Supernatants were analyzed using a Synapt XS Electrospray Ionization Quadrupole-Ion Mobility Separation Time-of-Flight (ESI-Q-IMS-TOF) mass spectrometer (Waters Corporation, RRID: SCR-026524) coupled to an Acquity Premier Ultra-Performance Liquid Chromatography (**UPLC**) system (Waters Corporation, RRID: SCR-027039). Samples were analyzed in both positive (capillary voltage: 1.5 kV) and negative (2.0 kV) electrospray ionization modes across a mass range of 50-1200 m/z. Lockmass calibration was achieved using a 200 pM leucine-enkephalin (Leu-Enk) solution (Waters, 186006013), introduced every 10 sec via a secondary orthogonal sprayer. The system operated in data-independent acquisition with elevated energy (MbSE) mode with a 0.25 sec acquisition time per scan and a collision energy ramp of 10–30 V.

Chromatographic separation was performed using a Premier Bridged Ethylene Hybrid (**BEH**) Amide column (100 × 2.7 mm, 1.7 µm pore size) (Waters, 186009508) fitted with a matching Vanguard guard column (5 mm length) (Waters, 186009510). The mobile phase consisted of *Buffer A* [0.1 % (*v*/*v*) aqueous formic acid (Fisher Scientific, A117-50)] and *Buffer B* [0.1% (*v*/*v*) formic acid in acetonitrile (Millipore Sigma, 900682)] at a constant flow rate of 400 µl·min^-1^. The gradient program was 5% *Buffer A* + 95% *Buffer B* (0-1 min), 95% *Buffer A* + 5% *Buffer B* (1-9 min), and 5% *Buffer A* + 95% *Buffer B* (9-11 min). A pooled quality control (**QC**) sample was prepared and run intermittently between and within batches to optimize for sample dilution, monitor instrument performance, and serve as a peak alignment reference. Injection volumes were 1 µl in positive mode and 2 µl in negative mode.

Data acquisition and initial review were conducted using MassLynx (v. 4.2). Files were imported into Progenesis QI (v. 3.0) for alignment, peak detection, and mass deconvolution, using standard parameters. Metabolite identification was first performed using the Metabolite and Tandem MS Database (**METLIN**) MS/MS library (2019). Remaining unidentified features were matched against the Human Metabolome Database (**HMDB**) using *in silico* fragmentation prediction (Wolf *et al*., 2010). A total of 3,258 distinct metabolite features were detected, of which 417 were annotated. The complete raw peak area dataset is provided in **Supplementary Table 4**. All downstream analyses were conducted using just annotated metabolites, unless otherwise stated. Metabolites common to or unique among groups were visualized using a Venn diagram (**Fig**. **3C**). Only metabolites detected in ≥ 50% of samples within a given group were included.

### Metabolomic analyses

Raw metabolite peak areas (**Supplementary Table 4**) were first normalized to the total RNA recovered in BEEO pellets following IOF isolation (**Supplementary Table 3**) to account for variation in organoid mass. Metabolites not detected in any given sample were assigned a value of 0 and, therefore, no data imputation was performed. Statistical analyses were subsequently conducted in MetaboAnalyst (v. 6.0). During initial data filtration, 40 features with constant or single values across all samples were identified and removed. Resulting peak areas were then log-transformed (base 10) and pareto-scaled (mean-centered and divided by the square root of the standard deviation of each variable).

PCA with 95% confidence intervals (**Fig**. **3D**) was performed using the permutational multivariate analysis of variance (**PERMANOVA**) function, with distributions based on Euclidean distance from the first two principal components. The sparse partial-least squares discriminant analysis (**sPLS**-**DA**) was performed using 5 components and 10 variables per component (**Fig**. **3E**). Model performance was evaluated using 5-fold cross-validations (**CV**) with an increasing number of components and a fixed 10 variables per component. To account for batch (animal) effects, differentially abundant metabolites were determined by linear modelling incorporating covariate adjustment (**Fig**. **3F**-**H**) using the *limma* linear regression approach and raw *P* ≤ 0.05 threshold (Ritchie *et al*., 2015; Pang *et al*., 2022). Boxplots of select corresponding metabolites were also exported and are presented in **Figure 2I**.

Metabolite set enrichment analysis (**MSEA**) was performed by first standardizing metabolite names against the HMDB, PubChem, and Kyoto Encyclopedia of Genes and Genomes (**KEGG**) databases; unrecognized metabolites were excluded. MSEA was then conducted using the *global test* (Goeman *et al*., 2004) against the Relational Database of Metabolomics Pathways (**RaMP**-**DB**), which integrates 3,694 curated pathways from KEGG, Reactome, and WikiPathways. Only metabolite sets containing ≥ 2 entries were included. Enrichment ratios were calculated as the ratio of observed (Statistic Q) to expected (Expected Q) metabolites within each pathway (Lu *et al*., 2022). Results were visualized using GraphPad Prism (v. 10.4.2, b.534) as multiple variable plots, with significance on the y-axis, enrichment ratio on the x-axis, and bubble diameter also proportional to the enrichment ratio (**Fig**. **2J**).

Biomarker analysis was also performed using MetaboAnalyst 6.0, leveraging the receiver operator characteristic (**ROC**) curve-based model evaluation function. Initially, raw peak intensities were filtered, normalized, transformed, and scaled as aforementioned. Seven metabolites were then manually selected for ROC analysis, which was conducted using the *Random Forests* algorithm. Specifically, 100 cross-validations were performed, with results averaged to generate a ROC curve, with 95% confidence intervals also presented (**Fig**. **2K**).

### In vitro embryo production

Slaughterhouse-derived cumulus-oocyte complexes (**COC**) were acquired from Simplot (Meridian, ID, USA) and shipped overnight post-collection. Transport occurred within an incubator (38.5 °C) using a specialized M199-based maturation medium. Approximately 22 h after initial COC harvest, *in vitro* fertilization (**IVF**) was conducted using a standard protocol (Simintiras *et al*., 2021a). In brief, initially COC underwent four sequential washes by manual serial transfer between wells (SPL, 30004) containing *BO-IVF Medium* (IVF Bioscience, 71004) to eliminate residual maturation medium and debris. Subsequently, groups of approximately 50 COC were placed into individual wells, each containing *BO-IVF medium*, and incubated at 38.5 °C and 5% CO_2_ under humidified air until sperm preparation was completed.

Conventional bovine semen straws from a single sire (ST Genetics, Navasota, TX, USA) were thawed by immersion in water at 37 °C for 45 sec. The contents were then expelled into a conical tube containing pre-equilibrated (37 °C) *BO-SemenPrep Medium* (IVF Bioscience, 71003). Spermatozoa were pelleted by centrifugation (300 × *g*, 5 min, RT) and resuspended in *BO-SemenPrep Medium*. This wash step was repeated, and the final pellet resuspended in *BO-IVF Medium* that had been pre-equilibrated overnight (38.5 °C and 5% CO_2_ under humidified air).

Sperm concentration was determined using a hemocytometer to achieve the addition of 2 × 10^6^ spermatozoa per 50 COC. Gametes were then co-cultured at 38.5 °C and 5% CO_2_ under humidified air for 18 h. Presumptive zygotes were denuded of cumulus cells by manual transfer into a 1.5 ml microcentrifuge tube containing pre-equilibrated (37 °C) *BO-Wash Medium* (IVF Bioscience, 51002) followed by gentle vortex for 5 min. Denuded zygotes underwent four sequential washes by manual serial transfer between wells containing *BO-Wash Medium*.

Groups of approximately 50 washed presumptive zygotes were transferred into individual wells with pre-equilibrated (37 °C) *BO-IVC Medium*. Culture proceeded for 7 days under light mineral oil (Fujifilm Irvine Scientific, 9305) at 38.5 °C under 5 % CO_2_ and 5 % O_2_ under 90 % N_2_.

### Embryo culture in IOF

On Day 7 post-fertilization, blastocysts were randomly allocated to one of four treatment drops (**Fig**. **4A**). Specifically, treatments consisted of (***a***) synthetic oviduct fluid (**SOF**) – unaltered commercial *BO-IVC Medium*, (***b***) PBS – 2.8% (*v*/*v*) sodium bicarbonate [7.5% (*w*/*v*) solution (Corning, 25035CI)] in 1× PBS, (***c***) IOF control – IOF extracted from vehicle control BEEO (**Fig**. **1A**) + 2.8% (*v*/*v*) sodium bicarbonate [7.5% (*w*/*v*) solution] in 1× PBS, and (***d***) IOF diestrus mimic – IOF from diestrus-mimic treated BEEO (**Fig**. **1A**) + 2.8% (*v*/*v*) sodium bicarbonate [7.5% (*w*/*v*) solution] in 1× PBS. Treatment drops were overlaid with light mineral oil and equilibrated for 2 h at 38.5 °C under 5 % CO_2_ and 5 % O_2_ under N_2_. A total of 428 blastocysts were distributed across the four groups and cultured over six independent replicates. Embryos were imaged by brightfield microscopy daily (**Fig**. **4B**) and collected on Day 10 post-fertilization. The number of visually alive (**Fig**. **4C**) and fraction of hatched blastocysts (**Fig**. **4D**) each day was observed and recorded.

**Figure 4.**
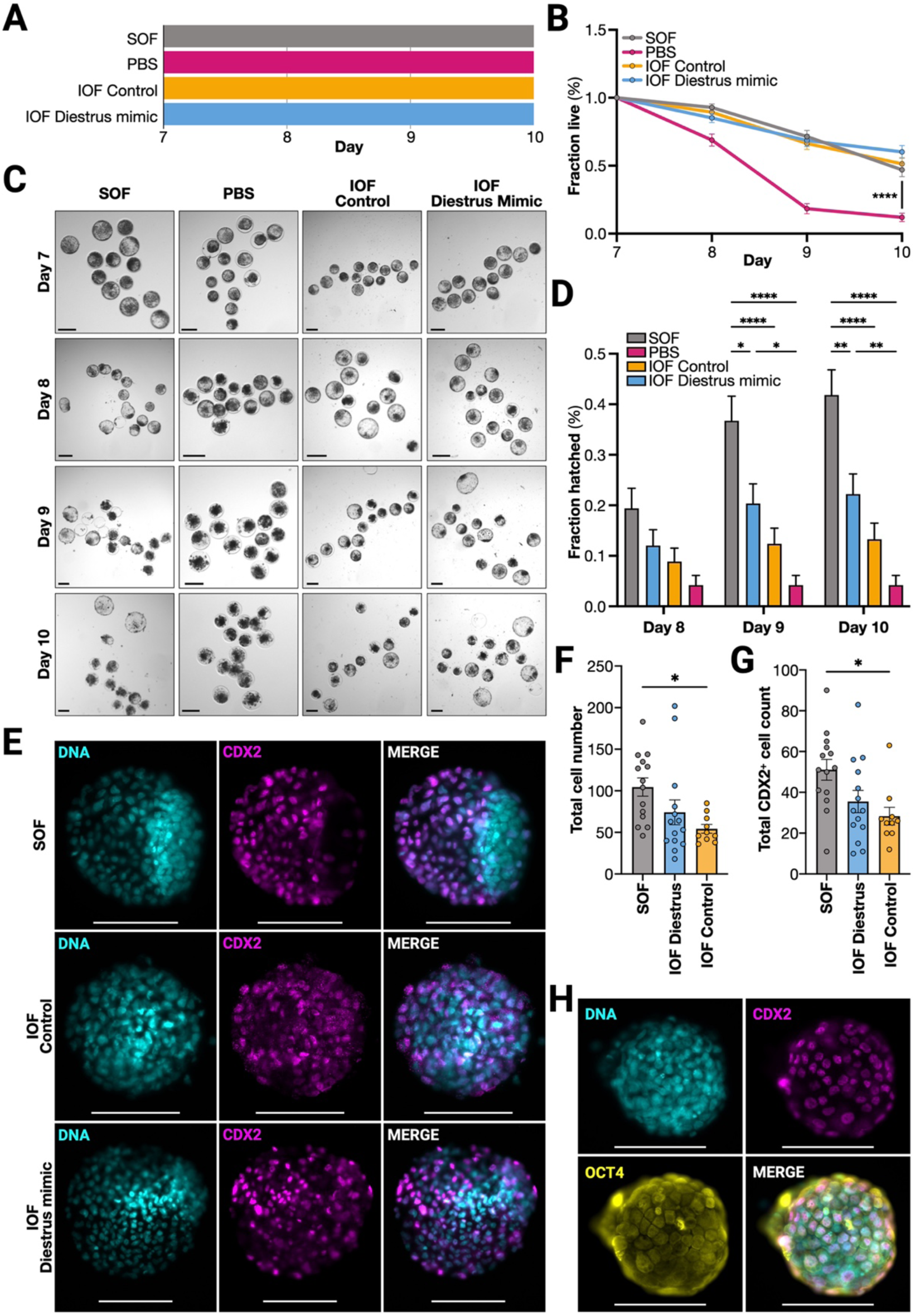
Bovine embryo culture in endometrial epithelial organoid (BEEO) intra-organoid fluid (IOF). (**A**) Dependent experimental variables – embryos were treated with a commercially available synthetic oviduct fluid (SOF), phosphate buffered saline (PBS), IOF from vehicle control treated BEEO, or IOF from diestrus mimic treated BEEO for 72 h. (**B**) Mean (±SEM) percentage of live embryos over culture duration. (**C**) Representative embryos according to treatment group and culture duration. (**D**) Mean (±SEM) percentage of embryos hatched according to treatment group and culture duration. (**E**) Representative embryo immunohistochemistry (CDX2) on Day 10 following each treatment. (**F**) Mean (±SEM) total embryo cell numbers on Day 10 following each treatment. (**G**) Mean (±SEM) CDX2 fluorescence intensity. (**H**) Representative embryo immunohistochemistry (CDX2 and OCT) on Day 10. All scale bars: 100 µm. Additional abbreviations: *P*≤0.05 (*), *P*≤0.01 (**), *P*≤0.001 (***), and *P*≤0.0001 (****).

### Embryo whole mount immunohistochemical labelling

On Day 10 post-fertilization all embryos underwent processing for immunohistochemical labelling, in accordance with an established protocol (Negrón-Pérez *et al*., 2017). In brief, embryos were sequentially washed by manual serial transfer between wells containing 1× PBS before fixation by immersion in 4 % (*v*/*v*) paraformaldehyde (Electron Microscopy Sciences, 15714) in 1× PBS for 20 min at RT. Following three additional sequential washes in 1× PBS, embryos were incubated with 0.5% (*v*/*v*) Triton X-100 in 1× PBS for 30 min, followed by a 1 h incubation in 5% (*w*/*v*) BSA in 1× PBS, at RT. Embryos were then incubated with the primary antibody (**Supplementary Table 2**) diluted to 1 µg·ml^-1^ in *Antibody Buffer* overnight at 4°C.

Thereafter, embryos were washed three times for 2 min each by transfer between wells containing 0.1 % (*w*/*v*) BSA with 0.1 % (*v*/*v*) Tween-20 in 1× PBS at RT. Embryos were then similarly incubated with the secondary antibody (**Supplementary Table 2**), diluted to 1 µg·ml^-1^ in *Antibody Buffer* for 1 h, at RT. This was followed by three additional washes using 0.1 % (*v*/*v*) Tween-20 with 0.1 % (*w*/*v*) BSA in 1× PBS for 2 min each, at RT. For staining with a second primary antibody, these steps were repeated. For nuclear counterstaining, embryos were transferred into Hoechst (1:5000 dilution in 1× PBS) for 5 min at RT, then washed three times in 1× PBS for 2 min each, at RT. Finally, embryos were transferred onto microscope slides using an EZ-Grip (Cooper Surgical, 7722802) and mounted using Fluoromount-G. Immunofluorescence imaging was performed as aforementioned (**Fig**. **4E**-**H**). Images were taken using a Leica DMI8 fluorescence microscope. Individual nuclei (**Fig**. **4F**) and CDX2^+^ (**Fig**. **4G**) nuclei were counted to determine total cell numbers using Fiji (ImageJ).

### Statistical analyses

Additional statistical analyses not previously described were performed using GraphPad Prism for Mac (v. 10.4.2, b. 534). More specifically, BEEO morphologies (**Fig**. **1C**-**E**) were compared by two-way analysis of variance (**ANOVA**) with multiple comparisons and subsequent Tukey’s *post hoc* test. IOF relative volumes (**Fig**. **3A**), embryo hatching proportions (**Fig**. **4D**), and cell counts (**Fig**. **4F**-**G**) were compared by ordinary one-way ANOVA, without matching or pairing, with multiple comparisons, and subsequent Tukey’s *post hoc* test. Asterisks denote significance as follows: *P*≤0.0001 (****); *P*≤0.001 (***); *P*≤0.01 (**); and *P*≤0.05 (*).

## RESULTS

### BEEO establishment and morphological characterization

Our first objective was to establish and morphologically characterize BEEO derived from estrous-synchronized cattle (n=4) under various hormonal treatments (**Fig**. **1A**). Following primary culture establishment, BEEO from all animals demonstrated cryopreservation capability with successful regeneration after several months of storage, confirming their biobanking potential. Brightfield imaging (**Fig**. **1B**) and morphometric analysis revealed no treatment effects on organoid count (**Fig**. **1C**), cross-sectional area (**Fig**. **1D**), or circularity (**Fig**. **1E**). However, BEEO cross-sectional area increased by Day 6 relative to Day 0 in all corresponding groups (**Fig**. **1D**). Similarly, immunostaining confirmed expression of canonical markers, including epithelial specific cytokeratin 18 (**CK18**), gland specific forkhead box A2 (**FOXA2**), estrogen receptors (**ESR1**, **ESR2**), and the progesterone receptor (**PGR**), similarly to *in vivo* (**Fig**. **1E**-**F**).

### BEEO transcriptomic response

Despite the absence of morphological changes of BEEO to treatment, RNA-seq revealed robust transcriptional responses to hormonal supplementation. PCA showed partial transcriptome separation by treatment (**Fig**. **2A**). Moreover, hierarchical clustering of top DEG identified six distinct functional clusters (**C1**-**6**) (**Fig**. **2B**), with GO enrichment implicating: (***C1***) extracellular matrix and cell junction disassembly, suppressed by P4; (***C2***) ion transport and import, suppressed by E2; (***C3***) antiviral defense, upregulated by E2+P4; (***C4***) immune response, induced by IFNτ; (***C5***) ciliogenesis, enhanced by E2; and (***C6***) cell cycle pathways, broadly downregulated with single-hormone supplementation (**Fig**. **2C**-**D**).

Direct comparisons of BEEO transcriptomes following supplementation identified 1,593 (E2 *vs*. MPA; **Fig**. **2E**), 2,783 (diestrus mimic *vs*. vehicle control; **Fig**. **2F**), 4,317 (pregnancy mimic *vs*. vehicle control; **Fig**. **2G**), and 476 (pregnancy mimic *vs*. diestrus-mimic; **Fig**. **2H**) DEG. A full list of DEG is provided in **Supplementary Table 6**. Furthermore, PGSEA confirmed and extended the finding that hormonal supplementation alters BEEO DEG enrichment for pathways related to metabolism and immune regulation, among others (**Fig**. **2I**).

### BEEO metabolomic response

IOF volume was unchanged across treatments (**Fig**. **3A**), and we confirmed protein presence in IOF, but not our PBS blank (**Fig**. **3B**), indicative of BEEO secretory capacity. A limited number of metabolites were uniquely detected between treatment groups (**Fig**. **3C**). PCA showed that IOF profiles were distinct from extra-organoid fluid and PBS controls, but separation between treatment groups was minimal (**Fig**. **3D**). Sparse partial least squares discriminant analysis (**sPLS**-**DA**) – a supervised method for resolving complex, non-linear variation (Lê Cao *et al*., 2011; Ruiz-Perez *et al*., 2020) – provided only marginally greater resolution (**Fig**. **3E**), demonstrating modest differences in IOF composition following hormonal treatment.

Although IOF metabolic profiles were broadly similar across treatments, select metabolites were differentially abundant. In the diestrus mimic *vs*. vehicle control comparison, 73 differentially abundant metabolites (**DAM**) were identified prior to covariate adjustment, including 2 annotated metabolites. After covariate adjustment, 44 DAM were detected, of which 14 were annotated (**Fig**. **3F**). Similarly, in the pregnancy mimic *vs*. vehicle control comparison, 108 DAM were identified prior to covariate adjustment, including three annotated metabolites. After covariate adjustment, 73 DAM were detected, of which 20 were annotated (**Fig**. **3G**). Moreover, in the diestrus *vs*. pregnancy mimic comparison, 42 DAM were identified prior to covariate adjustment, including 3 annotated metabolites. After covariate adjustment, 39 DAM were detected, of which 16 were annotated (**Fig**. **3H**). Further pathway enrichment analysis of IOF from pregnancy *vs*. diestrus mimic treated BEEO highlighted DAM related to arachidonic acid metabolism, lysine degradation, and the glucose-alanine cycle, among others (**Fig**. **3J**).

Given that statistically significant values often make poor biomarker predictors, and *vice versa* (Gränsbo *et al*., 2013; Lo *et al*., 2015), we used an established (Zou *et al*., 2007) machine learning algorithm (ROC analysis) to determine whether a panel of just seven metabolites could discern between IOF from pregnancy *vs*. diestrus mimic BEEO. The resulting area under the ROC curve (**AUROC**) value was 0.885 (**Fig**. **3K**), generally considered ‘good’ (Xia *et al*., 2013).

### Functional assessment of IOF on embryo development

To evaluate the functional capacity of BEEO secretions, blastocysts were cultured for three days (Days 7-10) in IOF (**Fig**. **4A**). This endpoint was selected based on preliminary data showing significantly decreased embryo survival beyond Day 10. The IOF used in this functional assay was extracted in PBS and buffered only with bicarbonate solution, containing no additional supplements, therefore directly reflecting the intrinsic secretory capacity of BEEO.

As expected, embryos cultured in PBS, buffered only with bicarbonate, did not survive to Day 10. In contrast, IOF from vehicle control and diestrus-mimic BEEO cultures supported embryo survival better than PBS and at levels comparable to commercial embryo culture medium, though without further improvement as hypothesized (**Fig**. **4B**). Representative images are shown in **Figure 4C**. Moreover, by Day 10, IOF from diestrus mimic treated BEEO supported blastocyst hatching better than PBS, but not IOF from vehicle control treated BEEO. However, hatching proportions in the diestrus mimic group remained below those observed in commercial culture medium (**Fig**. **4D**).

We next examined embryo morphology in terms of total and CDX2^+^ (trophectoderm marker) cell numbers (**Fig**. **4E**-**G**). Embryos cultured in IOF from vehicle control treated BEEO had fewer total cell numbers (**Fig**. **4F**) and cells expressing CDX2^+^ (**Fig**. **4G**) compared to commercial embryo culture medium. Therefore, by inference, lineage specification (trophectoderm to inner cell mass ratio) was unaffected by treatment. Finally, we observed that Day 10 embryos expressed OCT4 (pluripotency marker) throughout the trophectoderm and presumptive inner cell mass across all treatments (**Fig**. **4H**).

## DISCUSSION

The establishment of physiologically relevant *in vitro* models that accurately recapitulate endometrial function remains a critical challenge in reproductive biology. Here we demonstrate the successful establishment of BEEO that maintain hormonal responsiveness and secretory function, providing a powerful platform for investigating maternal-embryo communication *in vitro*. Our findings show that BEEO not only preserve an epithelial phenotype but also generate functionally relevant secretions capable of partially supporting embryo development, thus bridging the gap between traditional *in vivo* studies and traditional cell culture models.

First we show that BEEO can be cryopreserved and regenerated comparably to established organoid systems (Boers *et al*., 2016). Moreover, under the conditions described here, BEEO consistently maintained expression of key epithelial markers (CK18, FOXA2) and steroid hormone receptors (ESR1, ESR2, and PGR) throughout culture and hormonal treatments, accurately mirroring the *in vivo* epithelial phenotype. Moreover, BEEO showed no overt morphological changes in response to hormonal stimulation, which aligns with the *in vivo* bovine endometrium, where glandular architecture remains relatively stable across the estrous cycle (Wang *et al*., 2007), despite dramatic changes in transcriptomic (Bauersachs *et al*., 2005) and obviously secretory (Gray *et al*., 2001) activity.

Our transcriptomic analyses support the premise that BEEO preserve hormone-responsive functionality, contrasting with earlier reports of diminished endometrial gland function *in vitro* (Nishino *et al*., 2021). This could be due to their BEEO culture in significantly diluted hydrogel. Notably, the upregulation of *MSTN*, *ALDOA*, and *PDGFA* – genes encoding secretory proteins elevated in the pregnant endometrium on Day 13, coinciding with conceptus elongation initiation (Musavi *et al*., 2018) – specifically in diestrus mimic *vs*. control BEEO – suggests maintenance of P4 responsive secretory programs.

Similarly, pregnancy *vs*. diestrus mimic treated BEEO exhibited physiological immune modulation, including upregulation of several interferon stimulated genes, such as *CXCL17*, a major macrophage chemotactic factor (Burkhardt *et al*., 2014), and *CCL28*, which regulates epithelial chemotaxis (Mohan *et al*., 2017) – which parallels expression patterns in the bovine endometrium *in vivo* (Brewer *et al*., 2020). Furthermore, pregnancy simulation upregulated ISG-transactivating complex components *STAT1*, *STAT2*, and *IRF9*, mirroring profiles documented in the ovine endometrial glandular epithelium early during pregnancy (Choi *et al*., 2001). This transcriptional recapitulation validates BEEO as a physiologically relevant model of aspects of bovine endometrial physiology.

IOF metabolomic profiling revealed modest treatment-specific signatures, demonstrating some functional divergence in secretory output despite the absence of morphological changes. Moreover, an intriguing observation emerged when examining individual metabolite performance through ROC curve analysis. Notably, several metabolites achieved high area under the ROC curve (**AUROC**) values despite not reaching conventional statistical significance thresholds. AUROC values provide a measure of binary classification accuracy independent of significance cutoffs, with values generally classified as excellent (0.9-1.0), good (0.8-0.9), fair (0.7-0.8), poor (0.6-0.7), or fail (0.5-0.6) (Xia *et al*., 2013).

This metric offers complementary insights to traditional *P* values by assessing discriminatory power between conditions. For instance, isobutyrylcarnitine – previously shown to exhibit flux in bovine ULF (Tríbulo *et al*., 2019) – displayed an AUROC of 0.889 but did not achieve conventional statistical significance between diestrus and pregnancy stimulated IOF. To this end, metabolites with high AUROC values but marginal statistical significance may represent biologically meaningful markers that warrant further investigation.

To further test the physiological relevance of BEEO, we assessed the ability of IOF to support embryo development. Remarkably, IOF, despite being diluted approximately seven-fold in PBS, maintained embryo survival rates comparable to optimized commercial medium, and significantly exceeded PBS-only controls. This is particularly striking given that the IOF contained no supplemental growth factors, amino acid complexes, or serum replacements, directly reflecting the intrinsic secretory capacity of hormonally stimulated BEEO.

Furthermore, the observation that diestrus-stimulated IOF better supported trophectoderm specification and proliferation compared to control IOF supports the (***a***) importance of hormonal priming in creating a developmentally supportive ULF, and (***b***) efficacy of BEEO IOF as a relatively faithful ULF mimic, similarly to human EEO (Simintiras *et al*., 2021b). Nonetheless, using more concentrated IOF, extending exposure durations, using a higher P4 concentration – known to prime the endometrium for elongation *in vivo* (Clemente *et al*., 2009) are areas of current research that may enhance developmental support in future iterations.

Additional limitations and areas for improvement include increasing replication, scoring embryos by embryo competence index (Rabaglino and Hansen, 2024), and deriving BEEO from different stages of the estrous cycle. Here BEEO were derived from Day 5 post-estrus endometrium, potentially influencing their initial hormonal responsiveness. Derivation from different cycle stages might enhance functional capacity. It is also worth noting that we observed slower growth kinetics after approximately five passages, suggesting a finite expansion potential, though this requires further validation.

Moreover, while we demonstrate that IOF contains proteins that likely support embryo development, we did not perform comprehensive proteomic profiling. Such analysis would provide critical insights into the regulation and importance of specific growth factors, cytokines, carrier proteins, and enzymes that comprise the functional secretome *in vivo* (Forde *et al*., 2014; Gegenfurtner *et al*., 2020). Moreover, integrating proteomic data with our transcriptomic and metabolomic profiles would enable systems-level understanding of how hormonal stimulation coordinates multiple molecular layers to create a developmentally supportive microenvironment.

Finally, the absence of stromal and immune cell components in our model, while advantageous for isolating epithelial-specific responses, neglects important paracrine interactions that occur *in vivo*. Future development of more complex BEEO models incorporating multiple cell types, as in other systems (Rawlings *et al*., 2021; Gnecco *et al*., 2023; Shibata *et al*., 2024), would provide a more physiologically complete platform for studying maternal-embryo communication.

Despite these limitations, the current BEEO model represents a significant advance in our ability to study bovine endometrial epithelial function in a controlled, manipulatable system and using species-matched embryos. More specifically, additional future research initiatives may include the use of targeted (*e*.*g*., siRNA) interventions (Morgan *et al*., 2018), pharmacological inhibition, or metabolite supplementation (Zhang *et al*., 2022) to dissect mechanistic contributions of specific pathways in creating the supportive embryonic microenvironment.

In summary, we established and validated a BEEO model that mirrors key *in vivo* phenotypes. By generating IOF that serves as a functional analogue of uterine luminal fluid, BEEO provide a platform for elucidating cell-specific mechanisms governing early pregnancy establishment. This model bridges the complexity of traditional *in vivo* studies with the experimental control of *in vitro* systems, offering opportunities to dissect the maternal contributions to embryo development and identify factors critical for pregnancy success. As we continue to refine culture conditions and expand functional capabilities, the BEEO system promises to accelerate our understanding of maternal-embryo communication and potentially identify novel therapeutic targets for improving fertility outcomes in cattle and other livestock species.

## AUTHOR CONTRIBUTION STATEMENT

Conceptualization: ID, CAS. Experiments: ID, ZLB, DMF, YL, XZ, CRL, CAS. Data analysis: ID, SCL, AIVM, FD, XF, CAS. Data interpretation: ID, AM, KRB, JJB, PBA, CAS. Project guidance: AV, PHE, XF, KRB, JJB, CAS. Initial manuscript preparation: ID, CAS. Manuscript revision: ID, AM, JJB, CAS. Supervision: CAS. Funding acquisition: AV, XF, CAS.

## ACKNOWLEDGEMENTS

The authors thank Mr. Manuel “Boo” A. Persica in the Louisiana State University School of Animal Sciences and the staff at Coastal Plains Meat Company, Eunice, Louisiana, for their assistance with tissue collection. Supplementary Figure 1 was created in part using biorender.com. The authors also acknowledge the use of ChatGPT for improving sentence structure and readability in parts; however, the authors take full responsibility for the content of this manuscript. Moreover, aspects of these data were presented at the 2025 Society for the Study of Reproduction (SSR) meeting in Washington D.C.

## DATA AVAILABILITY

All data are included in the manuscript and/or corresponding Supplementary Material.

## FIGURE LEGENDS

**Supplementary Figure 1.**
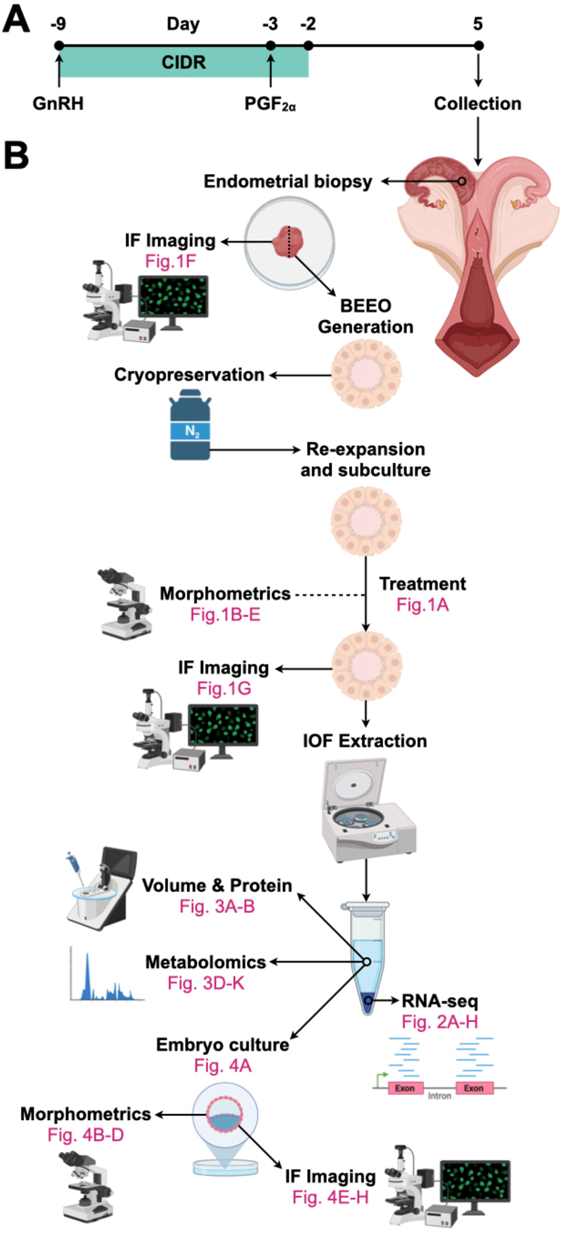
Schematic depiction of the study design including (**A**) animal synchronization protocol, and (**B**) experimental workflow. Abbreviations: gonadotropin-releasing hormone (GnRH), controlled internal drug release (CIDR) progesterone insert, prostaglandin F2⍺ (PGF2⍺), immunofluorescence (IF), bovine endometrial epithelial organoids (BEEO), intra-organoid fluid (IOF), and ribonucleic acid sequencing (RNA-seq).

**Supplementary Table 1**. List of culture and other media compositions used.

**Supplementary Table 2**. List of antibodies used.

**Supplementary Table 3**. Total organoid RNA recovered.

**Supplementary Table 4**. Raw RNA-sequencing counts

**Supplementary Table 5**. Raw metabolomic peak areas.

**Supplementary Table 6**. List of differentially expressed genes.

