## Supplementary figures and images for "Establishment and functional characterization of bovine endometrial epithelial organoids"

### Supplementary Figure 1

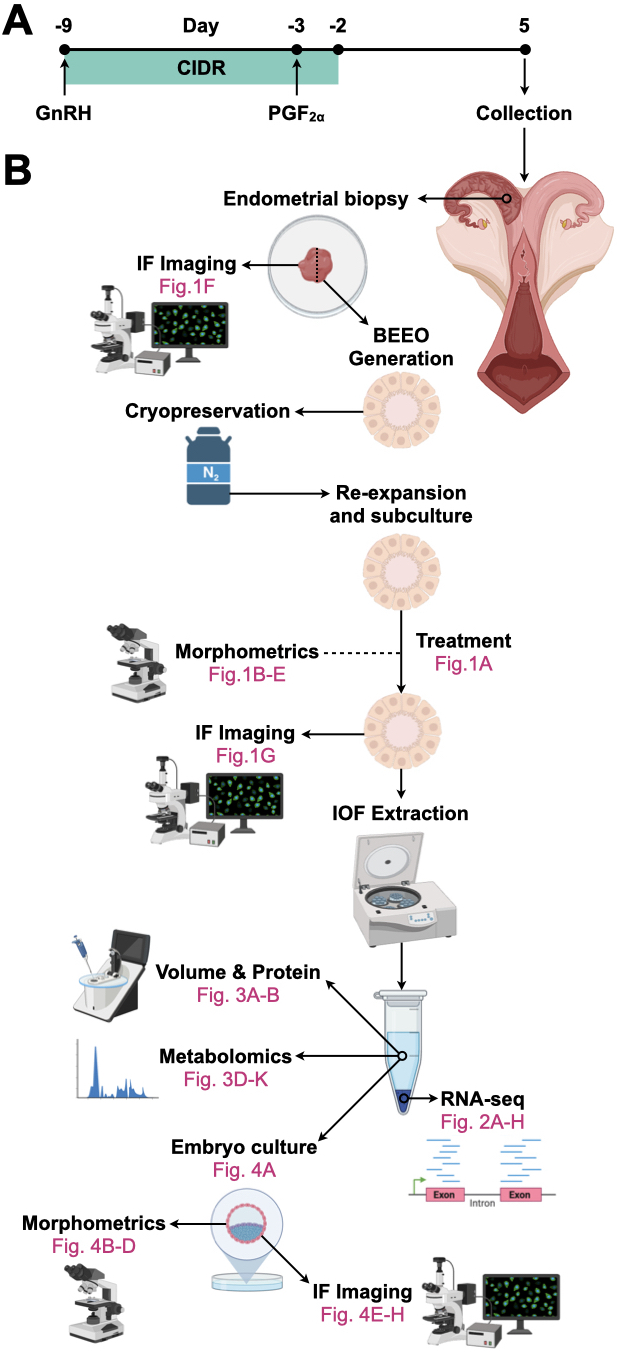
